# Dysregulation of Multiple Solute Carrier genes and Metabolic Deficits in *SLC1A4*-Mutant Human iPSC-Derived Hippocampal Neurons

**DOI:** 10.1101/2025.04.25.650669

**Authors:** Ritu Nayak, Omveer Sharma, Liron Mizrahi, Aviram Shemen, Utkarsh Tripathi, Yara Hussein, Wote Amelo Rike, Idan Rosh, Inna Radzishevsky, Hanna Mandel, Julia Ladewig, Tzipora C Falik Zaccai, Herman Wolosker, Shani Stern

## Abstract

Mutations in *SLC1A4*, which encodes the neuronal amino acid transporter ASCT1, disrupt metabolic and synaptic homeostasis, contributing to neurodevelopmental deficits commonly observed in autism spectrum disorder (ASD). To investigate the underlying molecular mechanisms of *SLC1A4*-related disorders, we utilized human iPSC-derived hippocampal neurons and applied an integrated multi-omics approach, combining electrophysiology, calcium imaging, metabolomics, proteomics, and transcriptomics. Our findings reveal an initial phase of early neuronal hyperexcitability, driven by increased sodium and potassium currents, followed by a progressive decline in synaptic activity at later stages. Metabolomic analysis identified elevated glycine, serine, and glutamate levels during early differentiation, contributing to excitotoxicity, whereas later glutamate depletion and extracellular matrix (ECM) disruption were associated with synaptic dysfunction. Proteomics data further showed dysregulation in metabolic pathways, amino acid biosynthesis, and fatty acid metabolism pathways during early time points, and in later stage dysregulation in metabolic and ECM-receptor interactions. Additionally, transcriptomic analysis revealed dysregulation in calcium signaling, amino acid metabolism pathways such as valine, leucine and isoleucine degradation, tryptophan metabolism, and glycine, serine, and threonine metabolism. Further investigation of SLC-family transporter genes uncovered disruptions in glutamate and glycine transport, establishing a direct link between amino acid transport dysfunction and neuronal deficits. Collectively, our study demonstrates that *SLC1A4* mutations lead to dysregulation of multiple solute carrier protein genes causing metabolic stress, excitability defects, and synaptic abnormalities, providing a molecular framework for understanding *SLC1A4*-related neurodevelopmental disorders and identifying potential therapeutic targets.

## 1. Introduction

Neurodevelopmental disorders (NDDs) represent a complex and diverse group of conditions that originate in childhood and involve disrupted brain development ^1^. Among the molecular pathways implicated in these disorders, solute carrier (SLC) transporters play a crucial role in maintaining ion balance, nutrient uptake, and amino acid transport across cellular membranes ^2^. SLC transporters, a superfamily of 439 members across 65 families, regulate the transport of ions, nutrients, and neurotransmitters across biological membranes ^3,4^. In the brain, 287 SLC genes have been identified ^5,6^. These transporters facilitate the exchange of glucose, amino acids, nucleic acids, ions, and other small polar molecules, playing a crucial role in maintaining metabolic homeostasis and cellular function in the brain ^7,8^. Within this family, the SLC1 subgroup encodes high-affinity glutamate transporters (EAATs) and neutral amino acid transporters (ASCTs), which are critical for synaptic plasticity and neuronal function ^9^.

One such gene, *SLC1A4* (solute carrier family 1 member 4) encodes for alanine/serine/cysteine/threonine transporter 1 (ASCT1) in the brain ^10^. ASCT1 is expressed widely throughout the body, with particularly high levels found in tissues like the brain, skeletal muscle, lungs, kidneys, ovaries, heart, and various parts of the digestive tract ^11^. This gene encodes for a protein that functions primarily as a sodium-dependent neutral amino acid transporter ^12^. In the brain, the *SLC1A4* gene plays a crucial role in the serine shuttle mechanism, which regulates L-serine and D-serine transport between astrocytes and neurons, which is essential for neurotransmission ^13^. Astrocytes synthesize L-serine via the PHGDH pathway, exporting it through ASCT1 (*SLC1A4*) to neurons, where it is converted into D-serine by serine racemase (SR) ^14^. Additionally, L-serine can be converted into glycine by the enzyme serine hydroxymethyltransferase, suggesting that the serine shuttle between astrocytes and neurons not only supports D-serine production but also indirectly contributes to glycine levels, both of which are essential for the proper functioning of NMDA receptors ^15,16^. This intricate shuttle system underscores the critical role of the *SLC1A4* gene in facilitating the metabolic interplay between astrocytes and neurons, which is essential for maintaining proper brain function ^18^.

Mutations associated with the *SLC1A4* gene have been reported to cause a rare autosomal recessive NDD, primarily characterized by global developmental delay, progressive microcephaly, spasticity, and seizures, often presenting in early infancy ^19,20^. First described by Srour and Damseh et al. in 2015, multiple pathogenic variants have since been identified across different populations ^20,21^. These mutations, including p.E256K, p.L315fs, p.R457W, and p.Trp453, disrupt ASCT1 function, impairing L-serine transport and neurotransmitter balance, leading to hypomyelination, cerebral atrophy, and severe neurological deficits^19,22,23^.

With the increasing identification of *SLC1A4* mutations, efforts have focused on developing mouse models to understand their neurodevelopmental impact and to explore potential treatments. Recent studies have introduced three mouse models: a full knockout (*SLC1A4*-KO), a knock-in model (*SLC1A4*-K256E) carrying the human p.E256K mutation, and a brain endothelial cell-specific knockout (*SLC1A4*tie2-cre) ^24^. These models revealed that *SLC1A4* is crucial for L-serine transport at the blood-brain barrier (BBB), and its loss leads to microcephaly, neurodegeneration, and synaptic dysfunction. Notably, early oral L-serine supplementation restored brain abnormalities, highlighting a potential therapeutic approach ^25^. Despite these advances, further research is needed to fully understand *SLC1A4*-related metabolic deficits and treatment strategies.

The advent of induced pluripotent stem cell (iPSC) technology has revolutionized the study of NDDs by enabling researchers to model disease-specific neuronal phenotypes using patient-derived cells ^26–35^. iPSC-based neuronal models provide a unique platform to investigate cellular and molecular mechanisms underlying NDDs, including mitochondrial dysfunction, hyperexcitability, and disruptions in calcium signalling. Studies on iPSC-derived neurons and oligodendrocytes from *SHANK3* mutation-associated Phelan-McDermid Syndrome ^31,34,26^ *UBTF* mutation-linked neurodevelopmental delay ^29^, and *DUP7* and other mutations associated with speech and cognitive disorders have demonstrated significant electrophysiological and synaptic alterations^29,30,36^ reinforcing the role of iPSC models in capturing patient-specific neuronal dysfunction. Given the challenges of directly studying patient brains, iPSC-derived neurons offer an ethically viable and biologically relevant approach to uncover disease mechanisms, identify biomarkers ^37,38^ and test potential therapeutic strategies. In this study, we have utilized an iPSC-derived model carrying the *SLC1A4* mutations and performed functional analysis of hippocampal neurons using patch clamp recordings and calcium imaging. Electrophysiological analysis demonstrated a hyperexcitability phenotype in early-stage *SLC1A4* mutant neurons, which showed a reduction in excitability at a later stage of maturation. To further investigate the molecular mechanisms underlying this phenotype, we explored a multi-omics approach, beginning with metabolomics to examine potential disruptions in amino acid metabolism. Our analysis revealed significant dysregulation of metabolites such as glutamate, glycine, and hydroxyproline during the early stages of neuronal maturation, which may underlie the early maturation and hyperexcitability phenotype observed during early time point. Interestingly, the abundance of these metabolites gradually declined at later stages of development. Further, proteomic and RNA sequencing analyses identified dysregulated pathways related to amino acid metabolism, synaptic signalling, and extracellular matrix dynamics. A theoretical explanation for these changes might be that *SLC1A4* mutations may affect other SLC genes, which in turn activate compensatory mechanisms that attempt to regulate these metabolites via alternative SLC transporters, leading to a broader disruption of amino acid and metabolic homeostasis, ultimately influencing neurophysiological function.

## 2. Materials and Methods

### 2.1. Ethics statement and blood collection

All participants, including patients and healthy controls, provided written informed consent for the study. The experimental procedures received ethical approval from the University of Haifa. Detailed information on the age and sex of all cell lines used in this study is summarized in the table below.

**Table 1:**
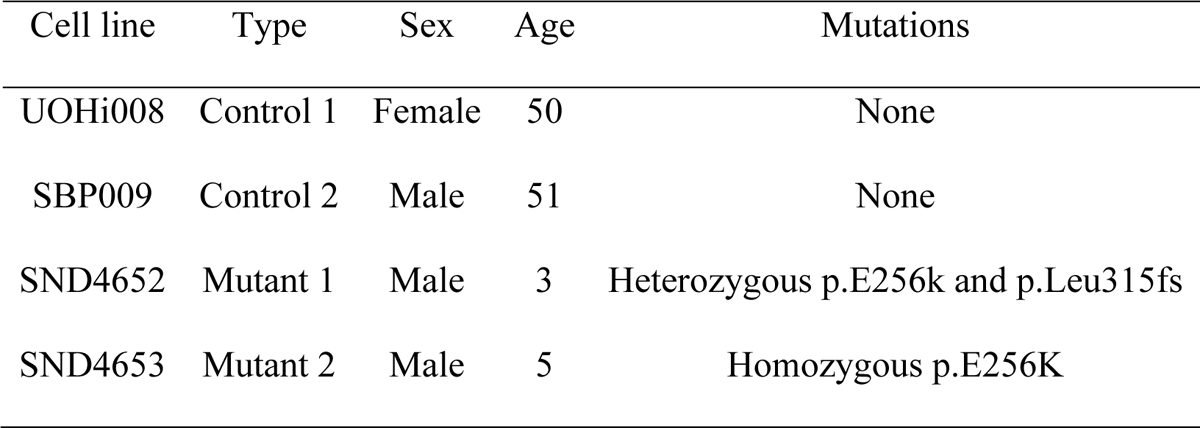
Details of *SLC1A4* patients and controls.

### 2.2. Isolation of peripheral blood mononuclear cells (PBMCs) from whole blood

PBMCs were isolated from whole blood collected in BD Vacutainer (Biogems). After collecting blood into BD vacutainers, the blood was processed immediately by centrifuging at 1800 ×g for 30 minutes at 23°C using a swing-out rotor. The plasma layer was carefully removed, and the PBMCs were collected. Cells were washed with DPBS without Ca²⁺ and Mg²⁺ and centrifuged at 300 ×g for 15 minutes. After removing the supernatant, cell were counted and resuspended in DPBS without Ca²⁺ and Mg²⁺. The PBMCs were then cryopreserved in FBS with 10% DMSO (Thermo Scientific, 127791000) at a concentration of approximately two million cells per cryovial. Cryovials were immediately placed in a Styrofoam container which was then immediately transferred to a −80°C refrigerator overnight and subsequently moved to liquid nitrogen for long-term storage.

### 2.3. Reprogramming of PBMCs into iPSCs using Sendai virus-mediated gene transfer

PBMCs were thawed four days prior to the day of reprogramming and cultured in StemPro™ SFM medium (Thermo Fisher Scientific, 10639011) along with SCF (Peprotech, Cat#300-07), FLT3 (Peprotech, Cat#300-19), IL3 (Peprotech, Cat#200-03) and IL6 (Peprotech, Cat#200-06) in a 24-well plate. The cells were fed daily by replacing 50% of the media. On the day of reprogramming (day 0), 500,000 cells were transduced with the CytoTune-iPS Sendai Reprogramming Kit (Life Technologies, A16518). Following transduction, the cells were centrifuged at 1,000xg for 30 minutes at 35°C and subsequently seeded in a 24-well plate, followed by overnight incubation at 37°C with 5% CO₂. The next day, the virus was removed by washing the cells, which were then plated into a low-attachment 12-well plate. After two days, the cells were transferred onto matrigel-coated plates for further culture, with media changes every two days. By day 7, the cells were gradually transitioned to ReproTeSR medium (Stem cell technologies, 05296), fully adapting to the new medium by day 8. Colonies began to emerge around days 15-21. Once the colonies were large enough, they were manually picked and transferred to new matrigel-coated plates and cultured in mTeSR plus medium (Stem cell technologies, 0274) for further expansion.

### 2.4. Differentiation of iPSCs into hippocampal neurons

Embryoid bodies (EBs) were generated by lifting the colonies using dispase (Stem Cell Technologies, 07923) for 15 minutes followed by mechanical dislodging of the iPSC colonies with a cell lifter. The dislodged colonies were transferred to low-adherence dishes and cultured in mTeSR medium. To initiate differentiation, the floating EBs were cultured in Dulbecco’s modified Eagle’s medium (DMEM)/F12, Invitrogen) supplemented with GlutaMAX (Thermo scientific, Cat#35050038), N2 (1:100, Gibco, Cat#17502048) and B27 without vitamin A (1:100, Rhenium, Cat#12587010). Additionally, an anti-caudalizing treatment cocktail was added, consisting of XAV939 (5 µM, Peprotech, Cat#2848932), SB431542 (10 µM, Biogems, Cat#301193), Cyclopamine (1 µM, Biogems, Cat#4445185), and LDN193189 Hydrochloride (0.1 µM, Biogems, 17504044) to induce differentiation for 20 days.

To obtain neural progenitor cells (NPCs), the EBs were plated after 20 days onto polyornithine/laminin-coated dishes (Sigma, Cat#P3655, Gibco, Cat#23017-015) in DMEM/F12 medium (Invitrogen) supplemented with GlutaMAX, N2, B27 without vitamin A, and laminin (1µg/ml) to allow neural rosette formation. After one-week, neural rosettes were manually collected based on morphology and dissociated using Accutase (AT-104, Innovative cell technologies). The dissociated cells were then replated onto polyornithine/laminin-coated dishes in NPC media, consisting of DMEM/F12, N2, B27 without vitamin A, laminin, and basic FGF (20ng/ml), to support NPC growth and proliferation.

NPCs were then plated on 6-well plates and cultured in DMEM/F12 medium supplemented with N2, B27 without vitamin A, L-Ascorbic acid (200 picoMol, Biogems, Cat #5088177), Dibutyl-cAMP (500 µg/ml, Adooq, Cat #A15914-5), laminin (1 mg/ml), brain-derived neurotrophic factor (BDNF) (20ng/ml, Peprotech, Cat #AF-450-02), and Wnt3a (20 ng/ml, R&D systems, Cat#5036) for two weeks. Following this period, the neurons were replated onto poly-L-ornithine/laminin coated coverslips, and the base media was switched to Brainphys (Stem Cell Technologies, Cat#05790) to promote neuronal maturation.

### 2.5. Immunocytochemistry

Coverslips containing iPSCs, NPCs, and hippocampal neurons were fixed for 15 minutes in 4% paraformaldehyde (PFA) pre-warmed to 37°C. After three washes with DPBS without Ca²⁺ and Mg²⁺, the cells were treated with a blocking and permeabilization solution composed of DPBS without Ca²⁺ and Mg²⁺, 0.2% Triton X-100, and 10% Donor Horse Serum for 60 minutes. The cells were then incubated overnight at 4°C with primary antibodies in the same blocking solution. iPSCs were stained for mouse anti-TRA-1-60 (Abcam, ab16288, 1:250), rabbit anti-OCT-4 (Abcam, ab19857, 1:250), mouse anti-SSEA4 (Abcam, ab16287, 1:250), and rabbit anti-NANOG (Abcam, 109250, 1:250). NPCs were stained for rabbit anti-PAX6 (CST, 60433, 1:250), and mouse anti-NESTIN (CST, 33475, 1:500). Neurons were stained for chicken anti-MAP2 (Abcam, ab92434, 1:500) and rabbit anti-PROX1 (CST, 14963, 1:250). Following overnight incubation, coverslips were rinsed three times with DPBS without Ca²⁺ and Mg²⁺(5 minutes each) and incubated for 60 minutes at room temperature with Alexa FluorTM secondary antibodies. A counterstain with DAPI (Abcam, ab228549, 1:3000) was then applied. The coverslips were washed, mounted onto slides using Fluoromount-G, and left to dry overnight in a dark environment. Fluorescence signals were captured using a Nikon A1-R confocal microscope, and images were further processed using NIS Elements 5.21 software and Imaris 9.8 for image analysis.

### 2.6. Electrophysiology

Whole-cell patch-clamp recordings were conducted in hippocampal neurons derived from *SLC1A4* patients as well as in hippocampal neurons derived from healthy controls. The recordings were done during two time points ∼5-9 weeks and ∼10-12 weeks post differentiation. The differentiation process was repeated 3-5 times. For each differentiation batch, recordings were performed on at least two coverslips, with data collected from ten neurons per coverslip. The culture coverslips were placed in a recording chamber filled with HEPES-based ACSF, composed of 139 mM NaCl, 10 mM HEPES, 4 mM KCl, 2 mM CaCl2, 10 mM D-glucose, and 1 mM MgCl2 (pH 7.5 and osmolarity adjusted to 310 mOsm), which was maintained at room temperature. Recording micropipettes, with a tip resistance of 10–15 MΩ, were filled with an internal solution containing 130 mM K-gluconate, 6 mM KCl, 4 mM NaCl, 10 mM Na-HEPES, 0.2 mM K-EGTA, 0.3 mM GTP, 2 mM Mg-ATP, 0.2 mM cAMP, 10 mM D-glucose, 0.15% biocytin, and 0.06% rhodamine (pH 7.5 and osmolarity adjusted to 290–300 mOsm). Data acquisition was performed at room temperature using Clampex v11.1, with a sampling rate of 20 kHz.

### 2.7. Electrophysiology analysis

For the study of hippocampal neurons, we conducted whole-cell patch-clamp recordings at two time points as mentioned earlier and the analysis were done accordingly.

#### 2.7.1. Total number of evoked action potentials

In the current-clamp mode, neurons were held at −60 mV with a constant holding current. Current injections were applied in 3 pA increments over 400 ms, starting 12 pA below the current required to maintain a −60 mV membrane potential. A total of 32 depolarization steps were performed. Neurons that required more than 50 pA to maintain the −60 mV potential were excluded from the analysis. The total number of action potentials was measured during the first 30 depolarization steps for hippocampal neurons.

#### 2.7.2. Analysis of sodium, fast and slow potassium currents

Sodium and potassium currents were recorded in voltage-clamp mode, with cells held at −60 mV and subjected to 400 ms voltage steps ranging from −90 to 80 mV. To account for differences in cell size, currents were normalized based on the cell capacitance, which was determined using a membrane test in Clampex software. The peak of the outward current, occurring within a few milliseconds of a depolarization step, was measured to assess the fast potassium current. In contrast, the slow potassium current was determined from the steady-state current at the end of the 400 ms depolarization period.

Sodium and potassium current amplitudes were analyzed statistically at specific voltage levels ranging from −20 to 0 mV for sodium currents and 40 to 80 mV for potassium currents. These voltage ranges were selected because they closely mimic the physiological conditions typical of a healthy neuron, providing relevant insights into the neuronal activity under normal functional states.

#### 2.7.3. Analysis of synaptic activity

In voltage-clamp mode, EPSCs were recorded to assess synaptic activity. Neurons were held at −60 mV while monitoring the currents in patched cells. Both the amplitude and frequency of synaptic events were analyzed using a custom MATLAB script. The cumulative distribution of EPSC amplitudes was calculated for each experimental group. To determine the rate of synaptic activity, the number of events was divided by the total recording time.

### 2.8. Calcium imaging experiments and analyses

Calcium imaging was conducted at two distinct time points on hippocampal neurons: the first at ∼5 weeks post-differentiation and the second at ∼9 weeks post-differentiation. To assess calcium transients, neuronal cultures were incubated for one hour in 2 mM Fluo-5 AM, which was dissolved in HEPES-based ACSF, composed of 139 mM NaCl, 10 mM HEPES, 4 mM KCl, 2 mM CaCl2, 10 mM D-glucose, and 1 mM MgCl2, adjusted to a pH of 7.4 and an osmolarity of 310 mOsm. Following the incubation, the culture coverslips were transferred into a recording chamber containing freshly prepared ACSF, pre-warmed to 37°C. Calcium imaging was then performed to observe the transients using a CCD digital camera and video microscopy (Leica Thunder imager). Data and images were collected and processed through time-lapse fluorescence microscopy, with each frame representing the fluorescence intensity at a given time point. Fluctuations in intensity indicated changes in intracellular calcium levels, reflecting neuronal activity. A custom MATLAB script was used to process the data in multiple stages.

First, a composite intensity map was generated by averaging pixel intensities across all frames, allowing for the identification of regions of interest (ROIs) corresponding to individual neurons. Candidate ROIs were selected by identifying fluorescence intensity maxima that exceeded a redundant detection. ROIs within 50 pixels of each other were removed. Additionally, pairwise correlation analysis of fluorescence intensity signals within selected ROIs was performed on highly correlated signals (r ≥ 0.9) which were considered likely to originate from the same neurons, leading to the removal of one of the redundant ROIs to maintain uniqueness.

The final ROIs were visually validated using the average intensity map integrating all frames. Calcium spike events were identified by detecting peak fluorescence intensity changes within the selected ROIs.To enhance signal clarity and minimize noise, low-pass filtering with a cutoff frequency of 0.01 Hz was applied to remove high-frequency noise, and fluorescence intensity signals were standardized to normalize variations across ROIs. Peaks were detected and subsequently fitted to an exponential decay model to characterize calcium transient dynamics. Key parameters extracted from the fluorescence intensity traces included rise time (Tr), representing the time taken to reach peak intensity from baseline; amplitude (ΔF/F0), indicating the maximum fluorescence change relative to baseline; decay time (Td), denoting the time required for the signal to return to baseline after reaching peak intensity; and decay constant (τ), describing the rate at which the fluorescence signal decreased post-peak. To ensure the robustness and reliability of calcium event identification, strict quality control criteria were applied. Thresholds based on adjusted R² values (statistical measure) were used to assess the goodness-of-fit of the exponential decay model, and calcium event duration constraints were implemented to eliminate artifacts and ensure physiologically relevant event detection. Computational tasks were parallelized using MATLAB’s parallel processing to speed up event detection and fitting. Finally, statistical analysis was performed to calculate the average event amplitude, event frequency (number of calcium events per unit time), and area under calcium events (summed fluorescence intensity relative to the baseline for each calcium event, reflecting the total calcium influx). Network bursts were identified using a sliding window approach to ensure the detection of sustained activity while reducing the influence of isolated spikes, where the cumulative neuronal activity was assessed within moving time windows. Each burst followed a characteristic pattern where the number of participating neurons increased, reached a peak, and then declined, forming a parabola-like shape. The sliding window method allowed for continuous monitoring of neuronal activity, capturing dynamic changes while smoothing out transient fluctuations. The burst was defined as the period from the initial rise in activity until the decline fell below a threshold (5% of neurons firing). We quantified burst features such as size (proportion of active neurons) and frequency (rate of burst occurrences over time).

To investigate the temporal characteristics of calcium imaging signals in control and *SLC1A4* mutant neurons, a detailed frequency domain analysis was performed. The fluorescence signals underwent pre-processing to eliminate baseline trends and artifacts, ensuring a clearer representation of the underlying signal dynamics. After pre-processing, the Fast Fourier Transform (FFT) was applied to the detrended signals to extract their frequency components. This transformation enabled the identification and quantification of dominant frequency components within the signals. The power spectral density was calculated to assess the relative contribution of each frequency component, providing deeper insights into the temporal fluctuations of calcium activity in both control and *SLC1A4* mutant neurons. The extracted frequency components were systematically compared between the two groups to examine potential variations in spectral distribution. The analysis specifically targeted fundamental and higher-order frequency components to evaluate differences in calcium signalling patterns. The results were visualized using spectral bar plots, highlighting the distinctions in frequency composition between both groups.

### 2.9. Mass spectrometry-based metabolomics

Hippocampal neurons derived from iPSCs were cultured in 12-well plates, with four replicates per cell line, each well containing approximately 1.5 million neurons. After ∼5-6 and ∼9-10 weeks post-differentiation, cells were washed gently with 1 ml of DPBS without Ca²⁺ and Mg²⁺ at room temperature. An ice-cold extraction buffer containing methanol: acetonitrile: water (5:3:2, v/v/v) was added to each well (200µl). The plate was then placed on a horizontal shaker for 5 minutes at a low RPM to facilitate extraction. The supernatant containing the extraction solution was collected into 1.5 ml eppendorf tubes and centrifuged at 16,000xg for 10 minutes at 4°C, after which the supernatant was immediately transferred to another tube, snap-frozen in dry ice, and instantly stored at −80°C for further analysis through liquid chromatography-mass spectrometry (LC-MS).

LC-MS analysis was performed at the Perlmutter Metabolomics Center, Technion following the protocol as mentioned by Mackay et al., 2015 ^39^. Briefly, a Thermo Vanquish Flex UPLC system coupled with an Orbitrap Exploris 240 Mass Spectrometer (Thermo Fisher Scientific) was used, with a resolution of 120,000 at 200 m/z, electrospray ionization, and polarity switching mode for both positive and negative ions across a 67–1000 m/z range. The UPLC setup included a ZIC-pHILIC column (SeQuant; 150 mm × 2.1 mm, 5 μm; Merck). Five microliters of biological extracts were injected, with compounds separated via a 15-minute mobile phase gradient, starting at 20% aqueous mobile phase of 20 mM (NH_4_)_2_CO_3_, pH 9.2 adjusted with 0.1% NH_4_OH (25%), and 80% acetonitrile, terminating at 20% acetonitrile. All metabolites were detected with mass accuracy below 5 ppm. Quality control samples, consisting of pooled aliquots, were analyzed to evaluate instrument variability, while samples were spiked with internal standards, including stable isotope-labeled glucose, glutamate, glutamine, lactate, alanine, and pyruvate. Blanks consisted of extraction buffer alone. Samples were normalized to volume (supernatant). Data acquisition was conducted using Thermo Xcalibur 4.4, and peak areas of metabolites were determined with Skyline 23.1.0.268 software. Metabolites were identified based on the exact mass of the singly charged ion and retention time was matched against an in-house library from commercial standards.

### 2.10. Metabolomics analysis

The raw data was normalized based on the protein concentration measured during metabolite extraction and processing. Following normalization, MetaboAnalyst 6.0 software was used for statistical analysis and visualization. This included generating the volcano plot, principal component analysis (PCA), and heat map (Fig. 4) to illustrate differential metabolite expression and patterns across different experimental conditions.

### 2.11. Sample preparation and quantitative proteomics

Proteomics analysis was performed at the Smoler Proteomics Centre, Technion, utilizing the LC-MS/MS facility. Briefly, the protein extraction was carried out using a modified protocol following the metabolomics procedure for both time points (∼5-6 and ∼9-10 weeks post-differentiation). The samples were lysed in 2% SDS and homogenized using syringes to facilitate protein denaturation and extraction. Then they were centrifuged at 12,000 RPM for 20 minutes at 4°C to separate the protein from cellular debris. The supernatant was collected and used for protein quantification, which was performed using the Pierce™ BCA Protein Assay Kit (Thermo Scientific, 23225).

For the LC-MS/MS analysis, the extracted proteins were analyzed using a Q Exactive HF mass spectrometer. Differentially expressed proteins were visualized through volcano plots generated using MATLAB scripts. To explore the biological significance of the identified differentially expressed proteins (DEPs), pathway analysis was conducted using the SHINYGO web-based tool.

### 2.12. RNA extraction and sequencing

Approximately 1.5 million hippocampal neurons were lysed using 800 µL of TRIzol™ reagent (Thermo Fisher Scientific, Cat no. 15596026). RNA purification was then performed using the Zymo RNA Clean & Concentrator kit, following the manufacturer’s instructions to ensure high-quality RNA isolation. The concentration and integrity of the extracted RNA were assessed using an ND-1000 Nanodrop spectrophotometer (Thermo Fisher Scientific) and a tape station. All samples exhibited RNA integrity numbers (RIN) above 7.

For RNA sequencing, libraries were prepared from neuronal RNA samples derived from *SLC1A4* patients and controls using the TruSeq RNA Library Prep Kit v2 (Illumina), in accordance with the manufacturer’s protocol. The quality of the raw sequencing reads (FASTQ files) was evaluated using FastQC (v0.11.5), and sequence alignment to the human genome (GRCh38.104) was performed using STAR (v2.7.9a). Differential gene expression analysis was carried out using DESeq2 (v1.34.0). The p-values were adjusted for multiple hypotheses using the Benjamini-Hochberg (BH) false discovery rate (FDR) correction. Genes were classified as differentially expressed if they had an adjusted p-value < 0.05 and an absolute log2 fold-change of at least 0.58. Based on these criteria, 2957 differentially expressed genes (DEGs) were identified in NPCs, while 4115 DEGs were detected in neurons. Following the identification of DEGs, pathway analysis was performed using the SHINYGO web-based tool.

## 3. Results

This section highlights the impact of SLC1A4 mutations on neuronal development and function. Electrophysiological recordings reveal accelerated maturation and hyperexcitability in immature neurons, along with reduced excitability and impaired synaptic transmission in mature neurons. These findings are further supported by multi-omics analyses.

### 3.1. Reprogramming and differentiation of *SLC1A4* patient and control samples

We investigated the expression of pluripotency and differentiation markers in iPSCs, NPCs, and differentiated hippocampal neurons derived from both control and *SLC1A4* mutant lines (control 1, control 2, mutant 1, and mutant 2 in Fig. 1). Immunofluorescence analysis of iPSCs revealed robust expression of key pluripotency markers, indicating that all iPSC lines maintained their undifferentiated, pluripotent state (Fig. 1a(i-iv) and b(i-iv)). Specifically, both control and *SLC1A4*-mutant iPSC colonies exhibited strong expression of SSEA4, a glycoprotein present on the surface of undifferentiated stem cells ^40^, and TRA-1-60, another surface marker indicative of pluripotency ^41^. In addition, the transcription factors NANOG and OCT4, critical for self-renewal^42^ and maintaining pluripotency ^43^, were also expressed in both control and mutant lines. DAPI staining confirmed nuclear integrity in all samples. Quantitative analysis of marker expression showed no significant differences between control and *SLC1A4* mutant iPSC lines, further confirming the maintenance of pluripotency in both groups (Fig. 1e(i-iv)).

**Figure 1:**
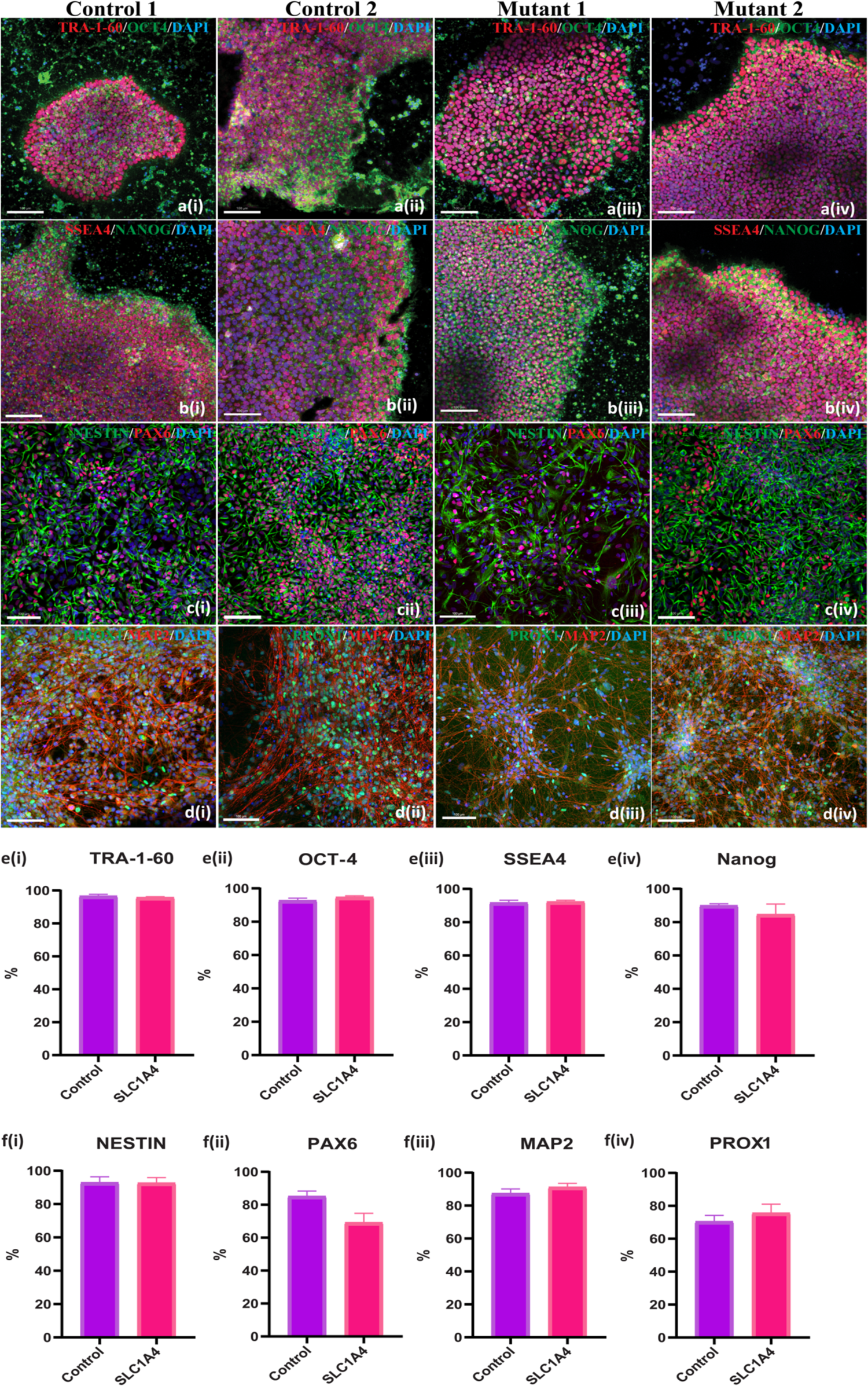
Characterization of pluripotency and neural differentiation in control and *SLC1A4* mutant and control lines. Immunofluorescence staining shows the expression of pluripotency markers TRA-1-60 (red) and OCT-4 (green) in iPSC colonies from control 1 (a.i), control 2 (a.ii), mutant 1 (a.iii), and mutant 2 (a.iv), with DAPI (blue) marking nuclei. Additional staining for SSEA4 (red) and NANOG (green) confirms pluripotency across control 1 (b.i), control 2 (b.ii), mutant 1 (b.iii), and mutant 2 (b.iv) lines. Differentiation into NPCs is indicated by NESTIN (green) and PAX6 (red) staining in control 1 (c.i), control 2 (c.ii), mutant 1 (c.iii), and mutant 2 (c.iv) NPCs. Further dentate gyrus granule neuronal differentiation is validated by MAP2 (red) and PROX1 (green) expression in control 1 (d.i), control 2 (d.ii), mutant 1 (d.iii), and mutant 2 (d.iv) neurons. Quantification of pluripotency markers shows comparable expression of TRA-1-60 (e.i), OCT4 (e.ii), SSEA4 (e.iii), and NANOG (e.iv) between control and *SLC1A4* mutant iPSCs. Similarly, quantification of differentiation markers indicates similar levels of NESTIN (f.i), PAX6 (f.ii), MAP2 (f.iii), and PROX1 (f.iv) between control and *SLC1A4* mutant-derived NPCs and neurons. Scale bars: 100 µm.

After the differentiation of iPSCs into NPCs (see Methods), immunocytochemistry staining for NESTIN and PAX6 was performed to check progenitor cell identity. Both control and *SLC1A4* mutant NPCs (control 1, control 2, mutant 1, and mutant 2 in Fig. 1) strongly expressed NESTIN and PAX6, markers of neural progenitors, indicating that the *SLC1A4* mutations did not impact NPC generation (Fig. 1c(i-iv)). Quantification showed no significant differences in the percentage of NESTIN-positive cells between control and *SLC1A4* mutant lines, confirming equivalent NPC formation (Fig. 1f(i)). While, PAX6, was expressed in all NPC lines, a slightly lower expression was observed in *SLC1A4* mutants compared to controls (Fig. 1f(ii)).

Neuronal differentiation was assessed by immunostaining for MAP2, a pan-neuronal marker, and PROX1, a marker for dentate gyrus granule neurons. Both control and mutant iPSC-derived neurons exhibited similar levels of MAP2 expression, demonstrating successful differentiation into neuronal phenotypes regardless of the *SLC1A4* mutations (Fig. 1d(i-iv)). Quantification confirmed no significant differences in the proportion of MAP2-positive neurons between control and *SLC1A4* mutant lines (Fig.1f(iii)). Similarly, no significant changes in PROX1 expression were observed, suggesting that the *SLC1A4* mutations did not disrupt hippocampal dentate gyrus granule neuron differentiation (Fig.1f(iv)).

### 3.2. Immature hippocampal neurons with *SLC1A4* mutations exhibit accelerated maturation and display a hyperexcitability phenotype compared to neurons derived from healthy controls

Using whole-cell patch-clamp recordings, we examined the electrophysiological properties of 79 control neurons and 183 neurons from patients with *SLC1A4* mutations. These neurons were derived from two control individuals and two *SLC1A4* patients and were recorded between 5-9 weeks post-differentiation. Figures 2a and 2b show representative traces of EPSCs in both control (a) and *SLC1A4* mutant neurons (b). Synaptic activity was measured by recording EPSCs at a holding potential of −60 mV. *SLC1A4* mutant neurons exhibited a significantly higher EPSC rate compared to controls (p = 0.0002, Fig. 2c), along with a notably increased mean EPSC amplitude (p < 0.0001, Fig. 2d). Furthermore, the cumulative distribution of EPSC amplitudes in *SLC1A4* mutant neurons displayed a rightward shift compared to control neurons, indicating larger EPSC amplitudes in the *SLC1A4* mutant population. This shift suggests an increased excitatory synaptic response in the *SLC1A4*-mutant neurons compared to controls (Fig.2e).

**Figure 2:**
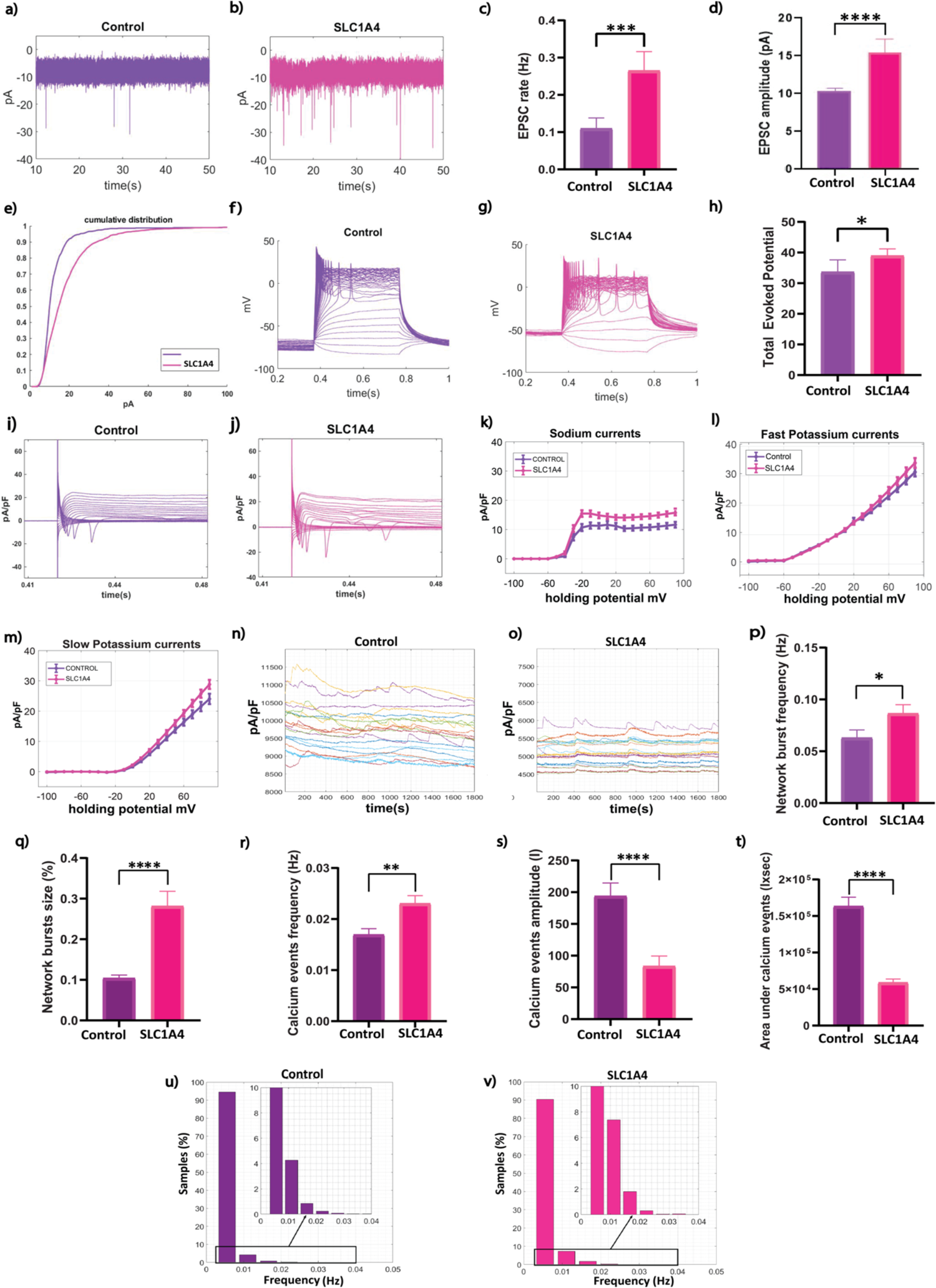
Electrophysiological and calcium imaging analysis of *SLC1A4* mutant and control neurons at 5-9 weeks post-differentiation. Representative traces of EPSCs in control (a) and *SLC1A4* mutant (b) neurons indicated enhanced synaptic activity in *SLC1A4* mutant neurons. Analysis of EPSC rate (p < 0.0002, c) and EPSC amplitude (p < 0.0001, d) revealed elevated synaptic activity in *SLC1A4* mutant neurons. The cumulative distribution of EPSC amplitudes showed a shift toward higher amplitudes in *SLC1A4* mutant neurons compared to controls (e). Representative traces of evoked action potentials in control (f) and *SLC1A4* mutant (g) neurons demonstrated increased excitability in *SLC1A4* mutant neurons. Quantification of the total number of evoked action potentials confirmed a significant increase in *SLC1A4* mutant neurons (p < 0.027, h). Representative traces of voltage-gated currents in control (i) and *SLC1A4* mutant (j) neurons were recorded. Quantification of sodium currents revealed a significant increase in *SLC1A4* mutant neurons, particularly between −20 and 0 mV (p = 1.83 × 10⁻⁵, k). Furthermore, differences were also observed in fast (l) and slow (m) potassium currents (p = 0.43 and p = 0.25, respectively). Representative traces of calcium transients in control (n) and *SLC1A4* mutant (o) neurons highlighted increased network activity in *SLC1A4* mutant neurons. Quantification of network burst frequency (p < 0.039, p) and burst size (p < 0.0001, q) showed significantly enhanced synchronous activity in *SLC1A4* mutant neurons. Event-wise analysis further revealed a significant increase in calcium event frequency (p = 0.0029, r), but a reduction in event amplitude (p < 0.0001, s) and area under the curve (p < 0.0001, t), indicating a higher rate of calcium events with reduced overall calcium influx per event. Frequency distribution histograms for control (u) and *SLC1A4* mutant (v) neurons supported these findings, demonstrating increased event frequency but reduced amplitude in *SLC1A4* mutant neurons. Data were collected from 3–5 independent differentiation cycles for each line. Statistical analyses were performed using one-way ANOVA for sodium and potassium current comparisons, and Mann–Whitney U tests for all other analyses. Statistical significance is indicated as follows: *p < 0.05, **p < 0.01, ***p < 0.001, ****p < 0.0001. Error bars represent the standard error of the mean (SEM).

Figures 2f and 2g show representative traces of evoked action potentials in control (f) and *SLC1A4* mutant neurons (g), further illustrating increased neuronal excitability in *SLC1A4* mutant neurons. Quantitative analysis revealed that the number of evoked action potentials was significantly higher in *SLC1A4* mutant neurons compared to control neurons (p = 0.027, Fig. 2h). Additionally, Fig. 2i, and 2j display representative recordings of voltage-gated sodium and potassium currents from control (i) and *SLC1A4* mutant neurons (j). Our analysis revealed a significant increase in sodium currents in *SLC1A4* mutant neurons, particularly between −20 and 0 mV, when compared to control neurons (p = 1.83 × 10⁻⁵, Fig. 2k). Furthermore, there were notable differences in both fast and slow potassium currents recorded between 40 and 80 mV in the *SLC1A4* mutant neurons compared to controls (p = 0.43, Fig. 2l, and p = 0.25, Fig.2m respectively). These findings were supported by one-way ANOVA analysis.

To further characterize functional activity, we examined calcium transients in neurons from control and *SLC1A4* mutant lines (see Methods). Representative traces of calcium transients from control and *SLC1A4* mutant neurons are shown in Fig. 2n, and 2o respectively. Quantification revealed significantly increased network burst frequency (p<0.039, Fig. 2p) and burst size (p<0.0001, Fig. 2q) in *SLC1A4* mutant neurons, indicating enhanced synchronized network activity. Additionally, the frequency of calcium events was significantly elevated in *SLC1A4* mutant neurons (p=0.0029, Fig. 2r). Whereas calcium event amplitude was significantly reduced (p < 0.0001, Fig. 2s). The area under calcium events was also significantly lower in *SLC1A4* mutant neurons compared to control neurons (p<0.0001, Fig. 2t), indicating a reduction in overall calcium influx per event. Spectral analysis of calcium activity at the early time point revealed that the fundamental frequency component (∼0.005 Hz) accounted for approximately 95% of the total spectral power in the control neurons and 91% in the *SLC1A4* mutant neurons, as shown in Fig. 2u and 2v. The second most dominant frequency component (∼0.011 Hz) constituted around 4.5% of the total power in control neurons, whereas in *SLC1A4* mutant neurons, this proportion increased to 7.5%. The third most component (∼0.016 Hz) was also more prominent in *SLC1A4* mutant neurons, suggesting altered frequency composition. These findings collectively indicate that *SLC1A4* mutant neurons exhibit a hyperexcitable phenotype during the early stages of maturation.

### 3.3. Mature hippocampal neurons with *SLC1A4* mutations exhibit reduced hyperexcitability and deficits in synaptic transmission compared to control neurons

The electrophysiological characteristics of 35 hippocampal neurons derived from two controls and 59 neurons from two patients carrying *SLC1A4* mutations were recorded using whole cell patch clamp ∼10-12 weeks post-differentiation. Figures 3a, and 3b present representative traces of EPSCs in both control (a) and *SLC1A4* mutant neurons (b). EPSCs were recorded at a holding potential of −60 mV. The EPSC rate was significantly lower in mutant neurons (p < 0.01, Fig.3c), while the EPSC amplitude was still higher (p < 0.05, Fig. 3d), suggesting reduced synaptic activity but larger synaptic events. A leftward shift in the cumulative distribution of EPSC was observed in mutant neurons, further indicating reduced synaptic activity (Fig. 3e).

**Figure 3:**
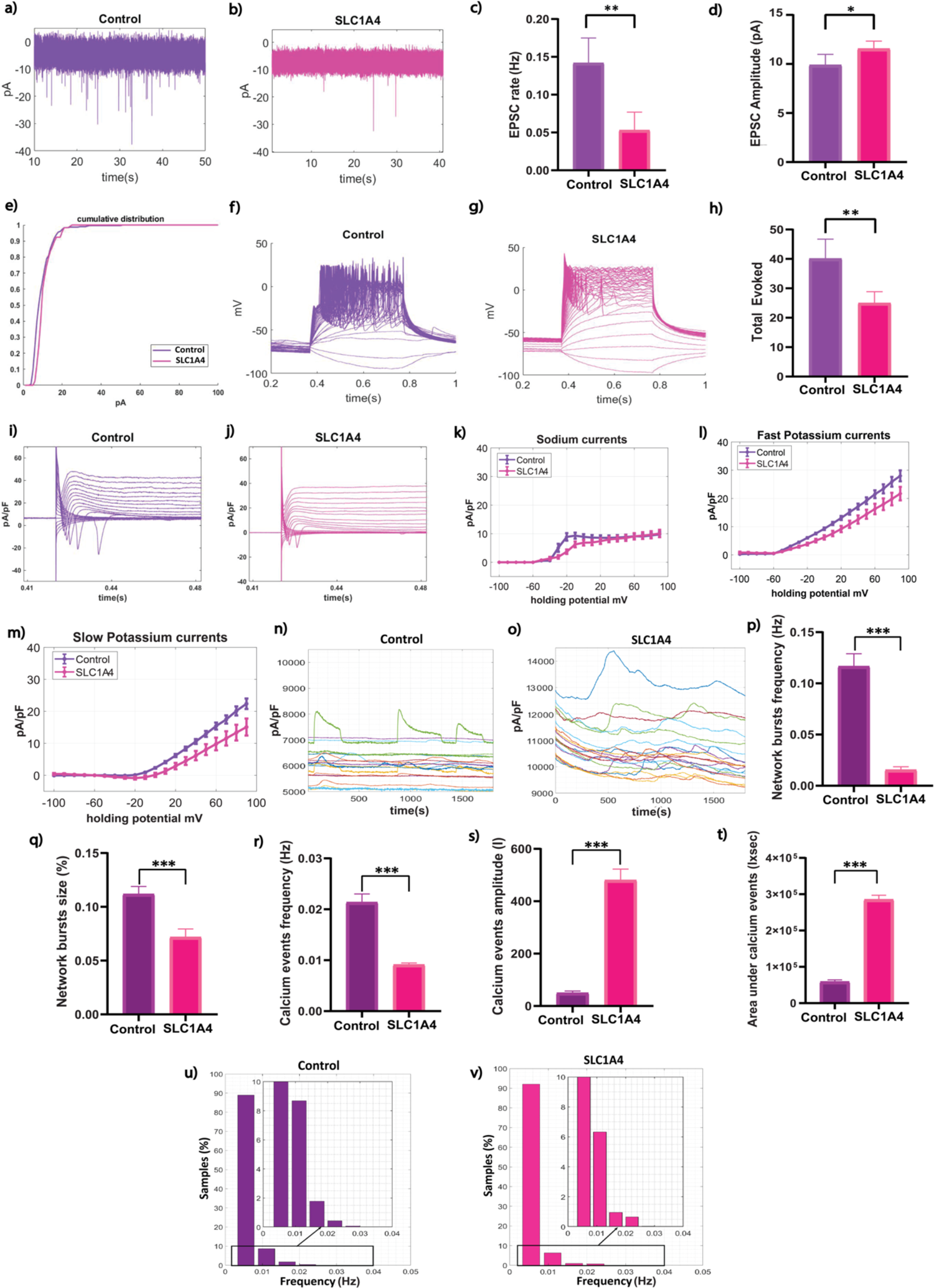
Electrophysiological and calcium imaging analysis of *SLC1A4* mutant and control neurons at 10-12 weeks post differentiation. Representative traces of EPSCs in control (a) and *SLC1A4* mutant (b) neurons indicated a decrease in synaptic activity. (c, d) Analysis of EPSC rate (p < 0.01, c) and EPSC amplitude (p < 0.05, d) revealed a lower frequency of synaptic events but a slight increase in amplitude in *SLC1A4* mutant neurons. (e) The cumulative distribution of EPSC amplitudes showed an overall shift in synaptic activity in *SLC1A4* mutant neurons. Representative current-clamp traces of evoked action potentials in control (f) and *SLC1A4* mutant (g) neurons demonstrated reduced neuronal excitability in *SLC1A4* mutant neurons. Quantification of total evoked action potentials showed a significant reduction in *SLC1A4* mutant neurons (p < 0.01, h). (i, j) Representative traces of voltage-gated sodium currents in control (i) and *SLC1A4* mutant (j) neurons were shown. Quantification revealed a significant reduction in sodium currents between −20 and 0 mV in *SLC1A4* mutant neurons compared to controls (p = 0.008, k). Differences in fast (p = 0.225, l) and slow (p = 0.019, m) potassium currents between 40 and 80 mV were also observed in *SLC1A4* mutant neurons compared to controls. Representative traces of spontaneous calcium activity in control (n) and *SLC1A4* mutant (o) neurons showed reduced network activity in *SLC1A4* mutant neurons. Quantification of network burst frequency (p < 0.0001, p) and burst size (p < 0.0004, q) confirmed significantly reduced network activity in *SLC1A4* mutant neurons. Quantification of calcium event frequency (p = 0.0029, r), amplitude (*p* < 0.0001, s), and area under the calcium transients (p < 0.0001, t) demonstrated fewer but larger calcium events in *SLC1A4* mutant neurons, suggesting altered calcium homeostasis. Frequency distribution histograms for control (u) and *SLC1A4* mutant (v) neurons showed a shift in network activity, with reduced event frequency and altered calcium dynamics in *SLC1A4* mutant neurons. Data were collected from 3–5 independent differentiation cycles per line. Statistical analyses were performed using one-way ANOVA for sodium and potassium currents and Mann–Whitney U tests for all other comparisons. Statistical significance is indicated as follows: *p < 0.05, **p < 0.01, ***p < 0.001, ****p < 0.0001. Error bars represent the standard error of the mean (SEM).

**Figure 4:**
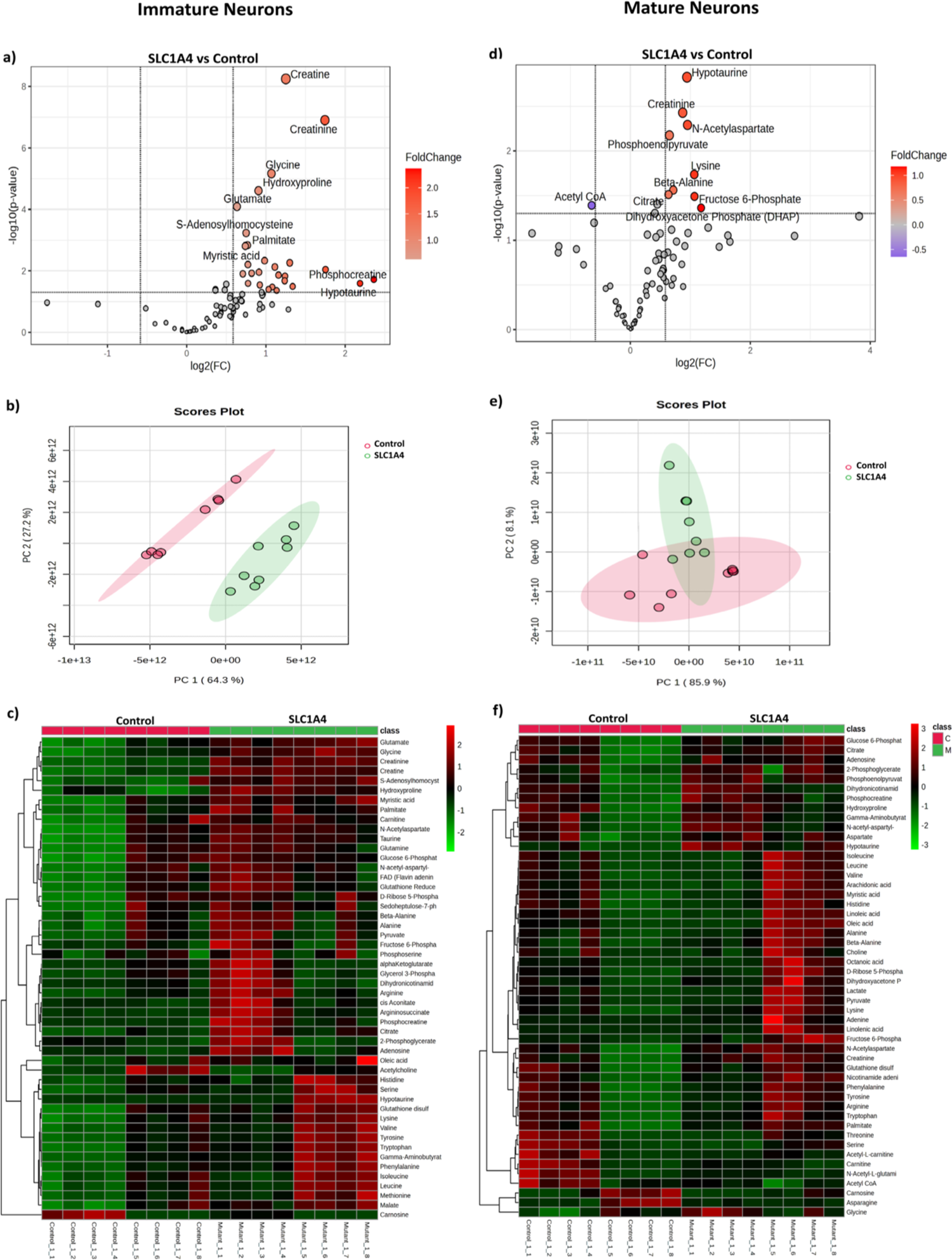
Metabolomic alterations in *SLC1A4* mutant neurons compared to healthy controls at different time points. (a–c) Immature neurons (∼5–6 weeks post-differentiation). (a) Volcano plots showed differentially abundant metabolites between *SLC1A4* mutant and control neurons. Each dot represented a metabolite, with red indicating significant upregulation and black indicating no significant change. Key upregulated metabolites in *SLC1A4* mutant neurons included creatine, creatinine, glutamate, glycine, serine, hydroxyproline, and phosphocreatine. (b) PCA plot demonstrated a clear separation between *SLC1A4* mutant and control neurons. (c) The heatmap illustrated the hierarchical clustering of 85 structurally annotated metabolites in control and *SLC1A4* mutant neurons. Red indicated upregulated and green indicated downregulated metabolites. (d–f) Mature neurons (∼9–10 weeks post-differentiation). (d) Volcano plot showed metabolite changes between control and *SLC1A4* mutant neurons. Upregulated metabolites in mutants included N-acetylaspartate, creatinine, phosphoenolpyruvate, and fructose 6-phosphate, while acetyl-CoA was significantly downregulated. (e) PCA plot showed partial separation between control and *SLC1A4* mutant neurons. (f) Heatmap displayed hierarchical clustering of 77 structurally annotated metabolites in mature neurons.

Representative traces of evoked action potentials for control (f) and *SLC1A4* mutant neurons (g) are shown in Fig. 3f and 3g, respectively. Quantitative analysis of evoked action potentials revealed that *SLC1A4* mutant neurons generated a significantly lower number of evoked action potentials compared to control neurons (p <0.01, Fig. 3h). Voltage-clamp recordings of sodium and potassium currents in control (i) and *SLC1A4* mutant neurons (j) are shown in Fig. 3i and 3j. A significant reduction in sodium currents was observed in *SLC1A4* mutant neurons (p = 0.008, Fig. 3k). Additionally, both fast and slow potassium currents showed differences between 40 and 80 mV with statistical significance for slow potassium currents (p = 0.22, Fig. 3l and p = 0.019, Fig. 3m respectively). One-way ANOVA supported these findings, indicating decreased ion channel activity in *SLC1A4* mutant neurons at a later time point.

In addition to patch clamp recording, calcium imaging revealed a marked reduction in network activity in *SLC1A4* mutant neurons. Representative traces of calcium transients from control and *SLC1A4* mutant neurons are shown in Fig. 3n, and 3o respectively. This reduction was reflected by a significant decrease in network burst frequency (p < 0.0001, Fig. 3p) and burst size (p < 0.0004, Fig. 3q). Calcium event frequency was also significantly reduced (p < 0.0029, Fig.3r), further supporting the observed loss of neuronal excitability over time. Interestingly, unlike the earlier time point, both calcium event amplitude (p < 0.0001, Fig.3s) and total area under calcium events (p < 0.0001, Fig.3t) remained significantly higher in *SLC1A4* mutant neurons, suggesting a shift in calcium homeostasis towards larger and prolonged calcium transients. At this later time point, the fundamental frequency component (∼0.005 Hz) accounted for approximately 89% of the total spectral power in control neurons and 92% in *SLC1A4* mutant neurons as shown in Figures 3u and 3v. The second most dominant frequency component (∼0.011 Hz) contributed ∼8.5% of the total power in the control neurons, whereas in the *SLC1A4* mutant neurons, this proportion decreased to 6.3%. The third most dominant frequency component (∼0.016 Hz) was observed to be more pronounced in the control neurons compared to *SLC1A4* mutant neurons. These results demonstrate that as *SLC1A4* mutant neurons mature, their synaptic properties change, including a reduction in hyperexcitability observed during the second time point.

### 3.4. The *SLC1A4* mutation induces metabolic alterations in immature and mature hippocampal neurons

To gain deeper insights into the molecular consequences of *SLC1A4* mutations and their associated neuronal dysfunction, we conducted LC-MS-based metabolomic profiling at two distinct time points. The first time point corresponded to immature neurons (∼5-6 weeks post-differentiation), where we compared *SLC1A4* mutant neurons to healthy controls. Differentiated cells were harvested, and metabolites were extracted for LC-MS analysis. Out of 85 structurally annotated metabolites, 27 were found to be significantly upregulated, and 57 were non-significantly changed in *SLC1A4* mutant neurons compared to controls (Fig. 4a). Among the upregulated metabolites were creatine, creatinine, and phosphocreatine, all linked to energy, metabolism ^44^. Glutamate, a primary excitatory neurotransmitter ^45^, along with glycine and serine, which are involved in neurotransmitter synthesis and signalling, were also elevated ^46^. Additionally, hydroxyproline, a product of collagen degradation, was increased, which may reflect collagen instability and disruption of extracellular matrix integrity_47._

To assess global metabolic changes associated with *SLC1A4* mutations, we performed PCA on metabolomics data (Fig. 4b). PC1 accounted for 64.3% of the total variance and separated *SLC1A4* mutant neurons from controls, indicating substantial mutation-driven metabolic divergence. PC2 explained an additional 27.2% of the variance. The clear separation observed in the PCA plot reflected consistent and significant metabolic alterations in *SLC1A4* mutant neurons compared to controls during the first time point. Furthermore, the heatmap further demonstrated the changes between the control and *SLC1A4* mutant groups. The heatmap complemented the volcano plot findings by illustrating the consistent upregulation of metabolites observed in immature neurons (Fig.4c).

We performed similar metabolomic profiling at second-time point (∼9-10 weeks post-differentiation) in mature neurons. Volcano plot analysis (Fig. 4d) revealed persistent but evolving metabolic disruptions. Differentiated neurons were collected, and metabolites were extracted for analysis. Of the 77 structurally annotated metabolites, 9 were significantly upregulated, 1 was downregulated and 66 metabolites were non-significant in *SLC1A4* mutant neurons compared to healthy controls. Volcano plot analysis highlighted the upregulation of several metabolites including creatinine, which remained elevated at the later time point. Notably, metabolites such as N-acetylaspartate, which plays a role in neuronal mitochondrial function was also upregulated ^48^. Additionally, metabolites involved in the glycolysis pathway, like phosphoenolpyruvate and fructose 6-phosphate, were also elevated, indicating altered glycolytic activity ^49^. Conversely, Acetyl CoA was significantly downregulated indicating a potential impairment in energy metabolism (Fig. 4d).

The PCA plot showed partial overlap between control and *SLC1A4* mutant iPSC-derived hippocampal neurons. PC1, accounts for 85.9% of the variance, reflecting substantial metabolic divergence driven by the mutations while PC2 captures 8.1% of the variance, showing some overlap, which suggests shared metabolic pathways among the groups (Fig. 4e). The heatmap (Fig. 4f) confirmed persistent dysregulation in amino acid, fatty acid metabolism and ECM related metabolites such as hydroxyproline, aspartate, choline, etc ^50^. These findings suggest that *SLC1A4* mutations induce early metabolic reprogramming, which is characterized by disrupted amino acid homeostasis, excitotoxic stress, and impaired energy metabolism among *SLC1A4* mutant neurons ^51^. This metabolic shift aligns well with our patch clamp and calcium imaging data which shows early stage neuronal hyperexcitability in immature neurons followed by reduced excitability as neurons mature.

### 3.5. Proteomic analysis points to metabolic dysregulation in *SLC1A4* mutant neurons compared to controls

We next carried out proteomic analysis to examine the impact of *SLC1A4* mutations in immature (∼5-6 weeks post-differentiation) and mature (∼9-10 weeks post-differentiation) hippocampal neurons. Fig. 5a Volcano plot displayed differentially expressed proteins (DEPs) in *SLC1A4*-mutant immature neurons compared to controls. Several proteins exhibited significant upregulation, such as SDSL(Serine Dehydratase-Like), linked to serine metabolism ^52^, and ACSF3 (Acyl-CoA Synthetase Family Member 3), a mitochondrial enzyme essential for fatty acid metabolism ^53^, CBR1(Carbonyl Reductase 1), a cytoprotective enzyme against oxidative stress ^54^. Furthermore, FPGT is involved in glycosylation ^55^. In contrast, DNALI1 (Dynein Axonemal Light Intermediate Chain 1) was downregulated, which may impair intracellular transport mechanisms essential for neuronal structure and function ^56^. Fig. 5b KEGG pathway analysis further identified dysregulated biological pathways in immature *SLC1A4*-mutant neurons. Notably, pathways related to metabolic pathways, amino acid biosynthesis, fatty acid metabolism, DNA replication, and cell cycle regulation were significantly altered. The expression of valine, leucine, and isoleucine metabolic enzymes was also notably disrupted, indicating dysregulated amino acid metabolism. Collectively, these findings highlighted early metabolic dysregulation associated with *SLC1A4*-mutant neurons, which further contributes to the hyperexcitability and synaptic alterations observed in *SLC1A4* mutant neurons in immature neurons.

**Figure 5:**
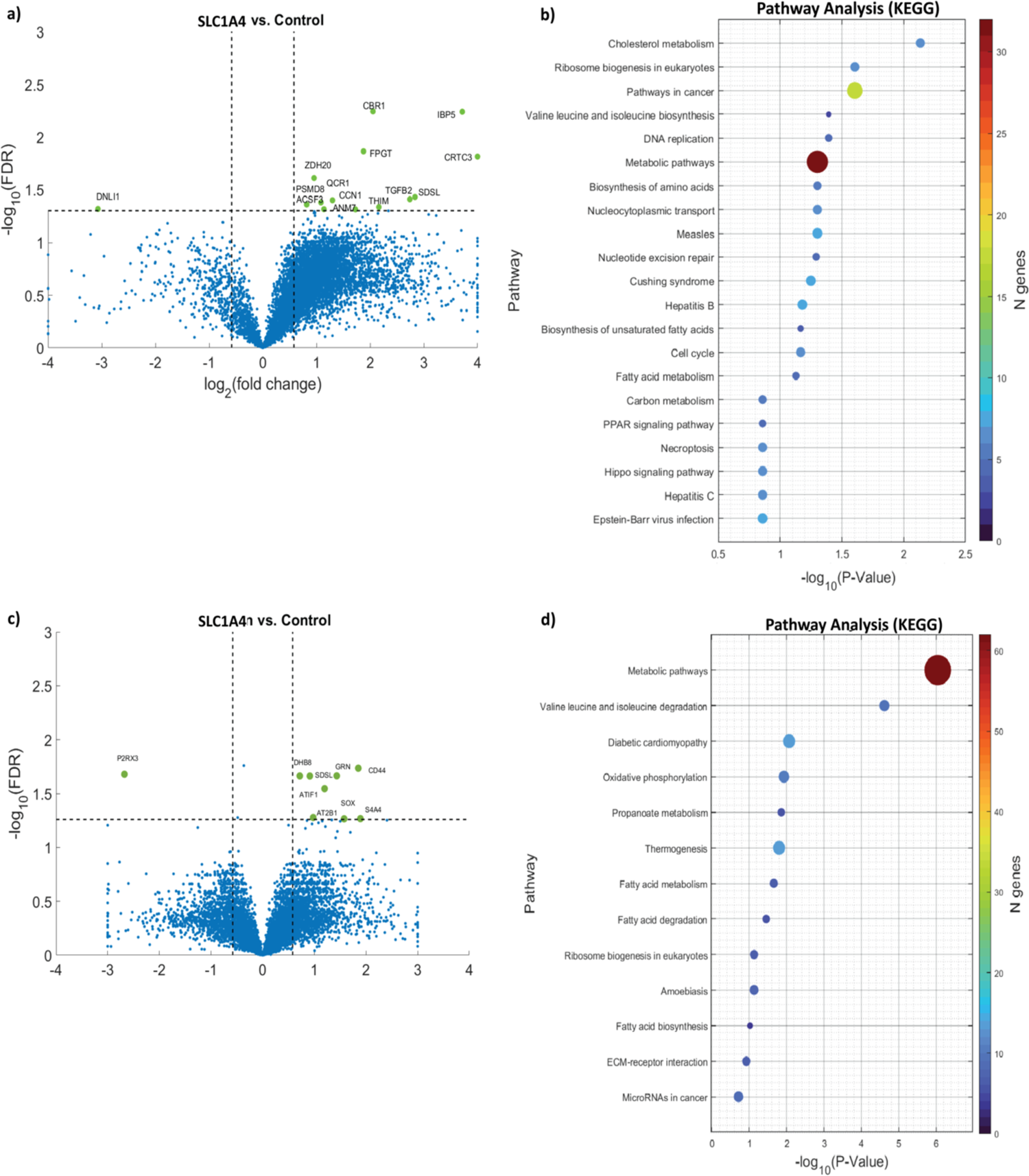
Proteomic analysis reveals metabolic dysregulation in *SLC1A4* mutant neurons. (a) A volcano plot depicting DEPs in immature neurons. (b) KEGG pathway enrichment analysis of DEPs in immature neurons, showing significant alterations in amino acid metabolism, and metabolic pathways. (c) A volcano plot of DEPs in mature *SLC1A4* neurons. (d) KEGG pathway enrichment analysis in mature neurons, indicating significant involvement of metabolic pathways, ECM-receptor interaction, amino acid degradation, and other key regulatory pathways.

In mature (∼9-10 weeks post-differentiation) neurons, the volcano plot (Fig.5c) revealed the upregulation of multiple proteins involved in cell signalling, metabolism, and structural regulation. SDSL, which was also upregulated in immature neurons, remained elevated, suggesting persistent dysregulation in serine metabolism. CD44, a cell surface glycoprotein involved in extracellular matrix interactions, was upregulated, potentially contributing to neuronal adhesion and structural remodeling^57^. SOX proteins, important transcription factors in neuronal differentiation and development, exhibited increased expression ^58^. Another protein ATP2B1, a plasma membrane calcium ATPase, was also found to be upregulated ^59^. In contrast, P2RX3, a purinergic receptor involved in synaptic function ^60^, was downregulated, potentially contributing to impaired synaptic transmission in mature neurons. KEGG Pathway analysis (Fig.5d) further highlights persistent metabolic dysregulation, Valine, leucine, and isoleucine, Fatty acid metabolism, and degradation pathway dysregulation. Notably, ECM-receptor interaction was enriched in mature neurons, indicating progressive alterations in extracellular matrix dynamics. Additionally, pathways linked to oxidative phosphorylation and propanoate metabolism were also found to be affected. Together, these findings underscore a shift from metabolic dysregulation to impairments in synaptic activity.

### 3.6. Solute carrier gene dysregulation in *SLC1A4* mutant NPCs and neurons

To investigate transcriptional alterations in *SLC1A4* mutant neurons, we performed RNA sequencing analysis of RNA extracted from NPCs and neurons (∼5-6 weeks post-differentiation). Fig.6a presents a heatmap of gene expression in *SLC1A4* mutant and control NPCs, illustrating differences between the groups. (Fig. 6b) Pathway analysis of differentially expressed genes (DEGs) in NPCs revealed multiple dysregulated pathways, including GABAergic synapse, calcium signalling, glutamatergic synapse, dopaminergic synapse, cholinergic synapse signalling, suggesting altered neuronal development and synaptic regulation. Although NPCs were not yet mature neurons and did not form functional synapses, pathway analyses revealed enrichment in synaptic-related pathways. This finding reflected the early transcriptional priming of NPCs toward neuronal lineages, where neurotransmitter-specific genes began to be expressed as part of fate specification. These pathways also exhibited important non-synaptic roles in neurodevelopment, influencing key processes such as NPC proliferation, migration, and differentiation. Similarly, Fig.6c shows a heatmap of gene expression in immature neurons (5-9 weeks post-differentiation), demonstrating differences between *SLC1A4* mutant and control neurons. Pathway enrichment analysis (Fig. 6d) highlighted key affected pathways in the neurons, including focal adhesion, ECM-receptor interaction, and axon guidance, suggesting potential disruptions in cell adhesion, signalling, and neuronal maturation. Pathways that relate to changes in metabolites such as metabolic pathways, valine, leucine, and isoleucine degradation, tryptophan metabolism, arginine and proline metabolism, propanoate metabolism, glycine serine and threonine metabolism, and more, suggest global dysregulation of metabolites caused by a single mutation in a transporter of alanine, serine, and cysteine. This may occur through homeostasis mechanisms trying to better regulate these metabolites through other SLC transporters and resulting in a global dysregulation of amino acids and metabolites.

**Figure 6.**
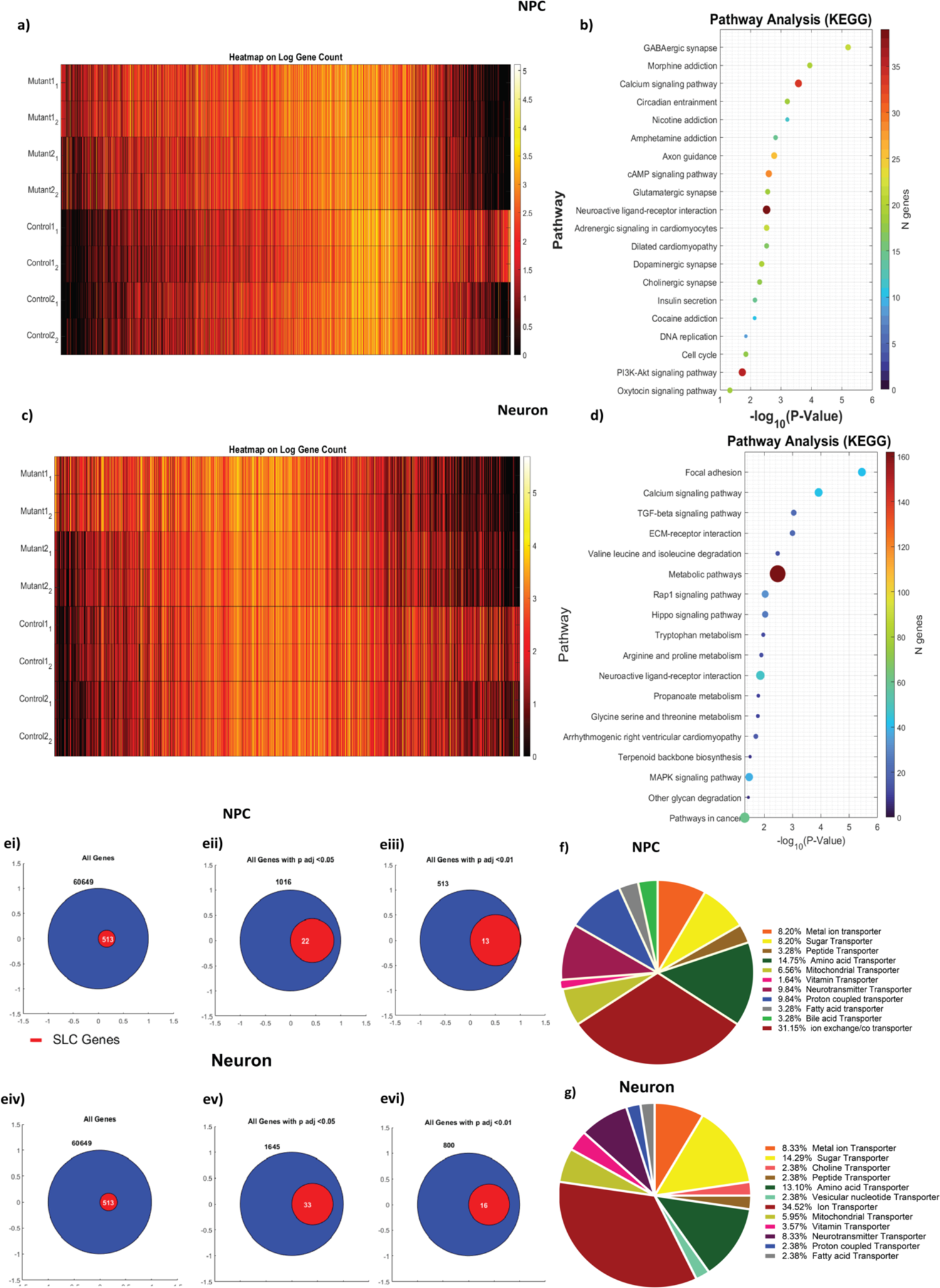
Solute carrier gene dysregulation in *SLC1A4* mutant NPCs and neurons. (a) A heatmap representing log-transformed gene expression levels in NPCs derived from control and *SLC1A4* mutant individuals. (b) KEGG pathway enrichment analysis for DEGs in NPCs, highlighting significantly affected biological pathways. (c) A heatmap showing log-transformed gene expression levels in iPSC-derived neurons from control and *SLC1A4* mutant individuals. (d) KEGG pathway enrichment analysis for DEGs in *SLC1A4* mutant neurons compared to healthy controls. (e(i) Venn diagram showing the total number of detected genes in NPCs, with a subset representing SLC family genes.(eii) Venn diagram displaying the number of differentially expressed genes (DEGs) in NPCs at an adjusted p-value threshold of <0.05, highlighting SLC genes among them. (eiii) Venn diagram illustrating the number of significantly dysregulated genes in NPCs at an adjusted p-value threshold of <0.01, showing SLC genes within this group.(eiv) Venn diagram representing the total number of detected genes in neurons, with a subset indicating SLC family genes.(ev) Venn diagram depicting the DEGs in neurons at an adjusted p-value threshold of <0.05, showing the proportion of SLC genes.(evi) Venn diagram highlighting significantly dysregulated genes in neurons at an adjusted p-value threshold of <0.01, emphasizing the presence of SLC genes in this subset.(f) A pie chart depicting the classification of SLC transporter-related genes in NPCs based on their functional categories. (g) A pie chart showing the classification of SLC transporter-related genes in neurons.

To further explore the impact of *SLC1A4* mutation on other solute carrier (SLC) transporters, Fig. 5e represents the proportion of SLC transporter genes within the total gene dataset for both NPCs and immature neurons. These included 513 transporter genes out of a total of 60,649 genes reported in the RNA sequencing data, accounting for approximately 0.85% of the genes, as shown in Fig. 6 (ei and eiv). However, 1,016 genes were dysregulated in NPC, including 22 SLC transporter-related genes (approximately 2.17%), as shown in Fig. 6 (eii). At a more stringent threshold of p < 0.01, 13 SLC genes remained significantly dysregulated as shown in Fig. 6(eiii). Similarly, in neurons, we identified 1,645 DEGs, among which 33 were SLC transporter genes (approximately 2.01%), as shown in Fig. 6 (ev). While 16 SLC genes remained significantly altered at p < 0.01 Fig. 6(evi), these findings indicate an increased proportion of SLC transporter genes among DEGs. The majority of dysregulated transporters in NPCs were classified under ion exchange transporters (31.15%), followed by peptide, amino acid, and mitochondrial transporters (Fig. 6f). In contrast, neurons exhibited a higher proportion of neurotransmitter transporters (8.33%) (Fig. 6g), suggesting a potential explanation for the synaptic deficits. These results reinforce the hypothesis that mutations in the *SLC1A4* gene may alter the expression of other SLC transporters essential for cellular transport mechanisms, potentially impacting neuronal function.

### 3.7. Multi-Omics pathway enrichment analysis of young neurons

Multi-omics pathway analysis revealed valine, leucine, and isoleucine biosynthesis and degradation pathway as the only pathway consistently dysregulated across metabolomics, proteomics, and transcriptomics datasets, highlighting a fundamental disruption in amino acid metabolism in *SLC1A4*-mutant neurons. Metabolomics analysis (Fig. 7a) further identified alterations in alanine, aspartate, and glutamate metabolism, glycine, serine, and threonine metabolism, and arginine biosynthesis, suggesting a widespread metabolic reprogramming affecting neurotransmitter balance and energy production. Proteomics data (Fig. 7b) showed dysregulation in the biosynthesis of amino acids, metabolic pathways, and amino acid metabolism pathways, indicating potential defects in protein synthesis and degradation. In transcriptomics significant changes were observed in calcium signaling and ECM-receptor interaction, pathways crucial for neuronal excitability, synaptic plasticity, and extracellular matrix stability (Fig. 7c). Based on this multi-omics pathway analysis, we gain a broader picture of the dysregulation across several amino acids which further contributes to dysregulation in their related pathways. These findings suggest that *SLC1A4* mutations might be responsible for the disruption in metabolic homeostasis and key signaling networks, leading to synaptic dysfunction and neuronal instability.

**Figure 7:**
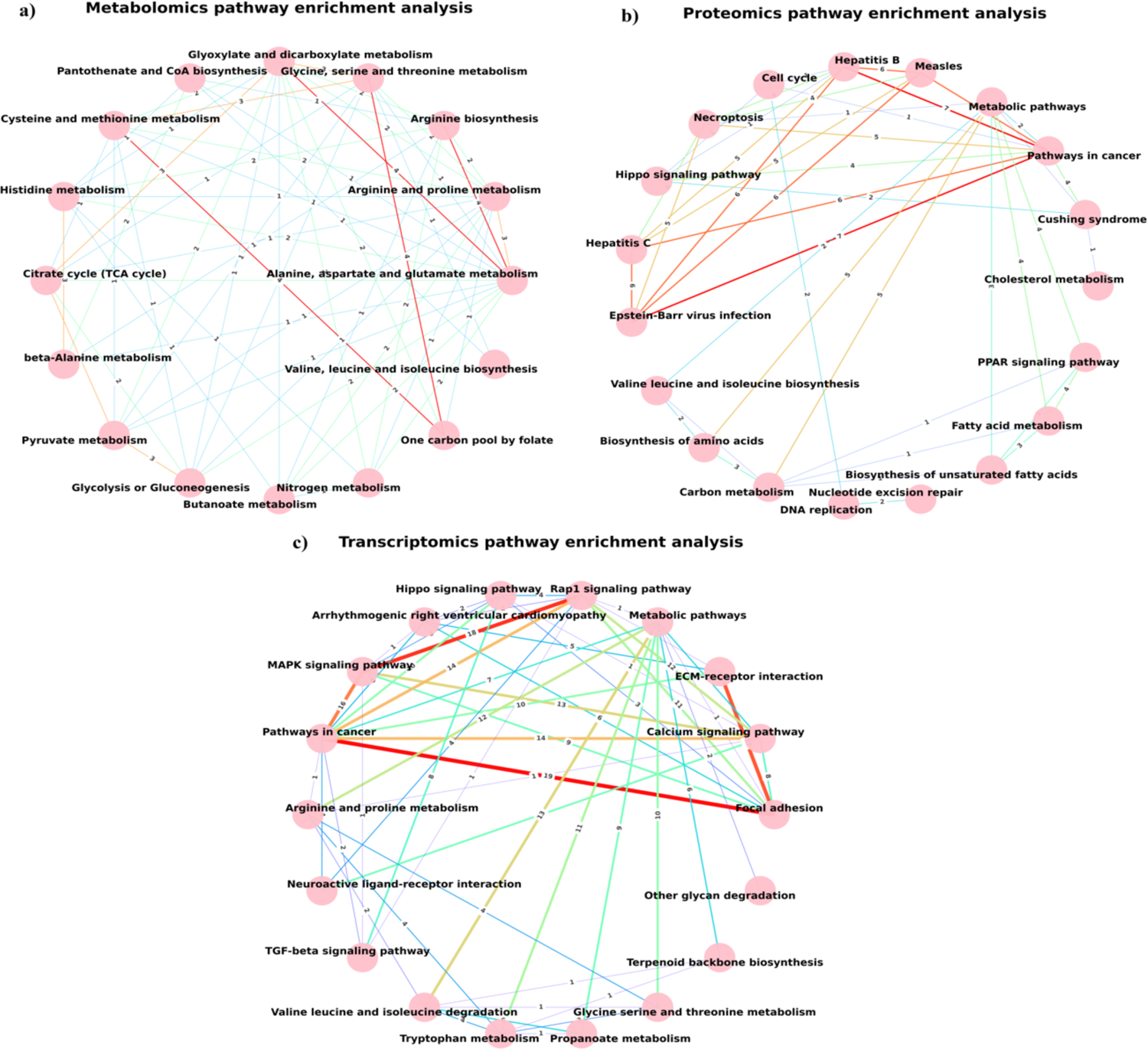
Pathway enrichment networks derived from (a) Metabolomics, (b) Proteomics, and (c) Transcriptomics data of young neurons. (a) Metabolomics pathway enrichment analysis reveals disruptions in amino acid metabolism, suggesting metabolic imbalances in *SLC1A4*-mutant neurons. (b) Proteomics pathway enrichment analysis highlights alterations in metabolic pathways, fatty acid metabolism, and calcium signalling (c) Transcriptomics pathway enrichment analysis shows significant dysregulation of pathways involved in calcium signalling, ECM-receptor interactions, and Rap1 signalling, which are crucial for neuronal excitability and synaptic plasticity.

## Discussion

This study provides a comprehensive multi-omics and functional characterisation into how pathogenic mutations in the *SLC1A4* gene disrupt neuronal function, contributing to a progressive, inherited metabolic disorder causing autism and a severe intellectual disability. By leveraging iPSC-derived hippocampal neurons, we characterized the distinct effects of *SLC1A4* mutation with a particular focus on neurons from the dentate gyrus (DG), a region where *SLC1A4* expression is highly enriched ^61^.

The successful differentiation of iPSC-derived hippocampal neurons was validated using a panel of pluripotency, NPC, and hippocampal-specific markers. This approach ensured the accurate characterization of neuronal populations. Immunofluorescence analysis confirmed that both control and *SLC1A4*-mutant iPSCs expressed key pluripotency markers, including TRA-1-60, OCT4, SSEA4, and NANOG, indicating robust reprogramming (Fig.1a-b,e). Upon neural induction, the expression of NESTIN and PAX6 was observed, demonstrating efficient NPC characterization (Fig. 1c, f). Further differentiation resulted in the expression of MAP2 and PROX1, with PROX1 being the well-established marker of hippocampal neurons in the DG, confirming the regional identity of the neurons in our neuronal population (Fig. 1d, f).

In our study, we first aimed to investigate the electrophysiological properties of *SLC1A4*-mutant neurons and uncover how these neurons behave over time. By combining electrophysiological recordings with calcium imaging, we observed a striking biphasic pattern in *SLC1A4* mutant neurons. During the early stage of differentiation, these neurons exhibited significantly enhanced sodium and potassium currents (Fig. 2k, 2l, 2m) and increased synaptic activity (Fig. 2c-e), suggesting an accelerated maturation in *SLC1A4* hippocampal neurons compared to healthy controls. Notably, our findings align with previous studies demonstrating premature excitability in iPSC-derived neurons derived from children with ASD, reinforcing the idea that accelerated maturation might be a shared feature across multiple neurodevelopmental disorders ^29,62,63,64^. However, upon maturation, *SLC1A4*-mutant neurons exhibited a notable reduction in excitability and synaptic transmission compared to controls (Fig. 3). By 11 weeks, they displayed significantly lower EPSC rate and network burst activity (Fig. 3c, p–q) and decreased calcium event frequency and amplitude (Fig. 3r–t).

These initial findings prompted us to further explore the underlying physiological alterations in *SLC1A4* mutant neurons. To deepen our understanding, we turned to a multi-omics approach, utilizing metabolomics, proteomics, and transcriptomics to gain insights into the molecular mechanisms associated with these changes. Given that *SLC1A4* encodes a solute carrier responsible for amino acid transport, we hypothesized that disruptions in this transporter might lead to widespread metabolic disturbances. Through metabolomics analysis, early in differentiation, glycine, hydroxyproline, serine, and glutamate were significantly elevated in mutant neurons (Fig. 4a). Glycine, an NMDA receptor co-agonist, accumulated more in *SLC1A4* mutant neurons, suggesting severe transporter dysfunction ^65,66^. Similarly, serine, a precursor of D-serine, was dysregulated, potentially contributing to early neuronal hyperexcitability (Fig.4a). Hydroxyproline, a product of collagen degradation and critical for extracellular matrix (ECM) stability, remained elevated across both time points, indicating continuous ECM dysregulation. In contrast, glutamate levels were significantly increased in the early phase, aligning with the heightened EPSCs and excitability observed in patch-clamp recordings (Fig.2f–h). These findings suggest that impaired function of *SLC1A4* likely disrupts glutamate homeostasis, leading to excitotoxic stress in young neurons ^67^. As neurons matured, excitability declined (Fig. 3), accompanied by a reduction in glutamate levels (Fig. 4d), indicating a progressive shift from hyperexcitability to synaptic dysfunction. This metabolic shift suggests that early excitatory overload may lead to long-term deficits in neurotransmission, ultimately contributing to synaptic abnormalities.

Building on our earlier findings, proteomic analysis of *SLC1A4* mutant and control neurons at two-time points revealed significant dysregulation in several proteins, consistent with the metabolic disturbances observed in metabolomics data. In immature neurons (∼5-6 weeks post-differentiation), volcano plot analysis identified a marked upregulation of SDSL (Serine Dehydratase-Like), a key enzyme in serine metabolism, as well as FPGT, a glycosylation-related protein (Fig.5a). Additionally, pathway analysis revealed disrupted biosynthesis of valine, leucine, and isoleucine, a finding that mirrored our metabolomics results for the first time point (Fig. 4a–c). Furthermore, the analysis highlighted enrichment in pathways associated with ribosome biosynthesis, cell cycle regulation, and DNA replication, suggesting disruption of protein synthesis (Fig.5b). These findings may point towards the early neuronal maturation, potentially underlying the early-stage hyperexcitability and synaptic alterations observed in *SLC1A4* mutant neurons(Fig. 2) ^68^.

As the neurons matured (∼9-10 weeks post-differentiation) proteomic analysis revealed that SDSL (Serine Dehydratase-Like) protein, which was upregulated in immature neurons, remained dysregulated and stayed elevated in mature neurons as well, suggesting persistent disruptions in serine metabolism (Fig.5c). Despite neuronal maturation, there remained persistent disruption of key metabolic pathways, including metabolic pathways, fatty acid metabolism, and degradation, valine, leucine, and isoleucine biosynthesis, as indicated by pathway analysis (Fig.5d). However, the downregulation of P2RX3, a purinergic receptor crucial for synaptic function, may contribute to impaired synaptic transmission in mature neurons. These impairments may be responsible for the long-term metabolic dysfunction that underlies the synaptic and functional deficits observed in mature neurons. Additionally, the enrichment of ECM-receptor interaction pathways in mature neurons further suggests progressive alterations in ECM dynamics, contributing to synaptic instability and dysfunction. When we further carried out transcriptomics analysis, we sought to investigate transcriptional changes in NPCs and immature neurons. Given that *SLC1A4* mutations exhibited a hyperexcitability phenotype during early time point, a characteristic often observed in ASD, we decided to focus our transcriptomic analysis on NPCs and immature neurons. In NPCs, cAMP signaling, calcium signaling, axon guidance, and PI3K-Akt signaling pathways were significantly altered, indicating early dysregulation in neuron formation and their connections. Moreover, we also found significant enrichment in neuron-specific pathways, including GABAergic, glutamatergic, dopaminergic, and cholinergic synapse signaling pathways indicating that NPCs are transcriptionally primed towards neuronal differentiation even in their undifferentiated state (Fig. 6a– b). In neurons, we identified persistent dysregulation in pathways related to valine, isoleucine, and metabolic pathways, which were consistent with the findings from our earlier omics analyses. These disruptions suggest a sustained alteration in amino acid metabolism that likely contributes to neuronal dysfunction. Additionally, we observed significant alterations in other metabolites such as tryptophan, arginine, and proline metabolism pathways, as well as propionate metabolism and glycine, serine, and threonine metabolism pathways (Fig. 6d). Alteration in these pathways further support the idea that metabolic reprogramming in *SLC1A4* mutant neurons may be associated with these key metabolic pathways. Furthermore, calcium signaling was notably dysregulated. These pathways have been previously associated with ASD, where disruptions in calcium homeostasis and neurotransmitter signaling contribute to synaptic dysfunction, neuronal hyperexcitability, and imbalances in excitatory-inhibitory transmission ^69,70,71^. Additionally, ECM-receptor interaction and Rap1 signaling also emerged as significantly affected pathways, consistent with the persistent elevation of hydroxyproline as observed in metabolomics (Fig. 4) and proteomics data (Fig. 5), thus suggesting progressive structural instability, with ECM-related pathways which also has been previously linked to ASD etiology ^72,73,74^.

Furthermore, our transcriptomic analysis identified significant alterations in SLC family genes in both NPCs and neurons (Fig. 6f–g). Therefore, we focused on transporter-related gene expression and observed significant alterations in genes associated with sodium ion exchangers, amino acid transporters, and neurotransmitter transporters. This disruption, along with the involvement of SLC transporters, led us to hypothesize that *SLC1A4* mutations might be responsible for triggering alterations in other SLC transporters as a compensatory mechanism that attempts to regulate metabolites via alternative transporters. This results in broader disruptions of amino acid and metabolic homeostasis, which ultimately affects neurophysiological function. Together, these findings provide a comprehensive view, illustrating how early metabolic disturbances initiate a cascade of molecular events that ultimately lead to synaptic dysfunction and structural instability.

This integrated analysis provides valuable insights into the biphasic nature of *SLC1A4*-associated pathologies, where initial metabolic dysregulation primes neurons for early hyperexcitability, followed by progressive synaptic dysfunction. By elucidating the molecular events associated with these changes, our findings offer a clearer understanding of the underlying mechanisms of *SLC1A4* mutations. This knowledge not only deepens our comprehension of the mutation’s impact on neuronal function but also opens avenues for developing targeted therapeutic approaches.

## Acknowledgements

The graphical images for this publication were plotted in MATLAB and MetaboAnalyst version 5.0. The authors would like to thank Ms. Bella Agranovich and Ifat Abramovich for expert guidance in metabolomics experiments, Dr. Sandra Horschitz (Hector Institute) for generating IPSCs, and Dr. Limor Kalfon (Galilee Medical Center, Israel) for obtaining patient blood samples.

## Declarations

Funding: - The Zuckerman STEM leadership program and Israel Science Foundation grants - 1994/21 and 3252 /21 to Prof. Shani Stern. Prof. Herman Wolosker was funded by the Laura and Isaac Perlmutter Foundation, Allen and Jewel Prince Center for Neurodegenerative Disorders, Israel Science Foundation (#279/24), Rappaport Family Institute for Research in the Medical Sciences, National Institute of Health/ National Institute of Neurological Disorders and Stroke (R01NS098740).

## Ethics approval and consent to participate

All participants provided informed consent before participating in the study. Approval for the study was obtained from the Research Ethics Board of the University of Haifa, Israel.

## Author contribution

Ritu Nayak (RN) and Omveer Sharma (OS) (co-first author) contributed equally to this work. RN grew the cells, conducted experiments, performed data analysis, drafted, and compiled the manuscript. OS conducted experiments, performed the data analysis, drafted, and compiled the manuscript. Liron Mizrahi (LM) performed experiments and Aviram Shemen (AS) performed experiments and assisted with data analysis. Utkarsh Tripathi (UT) assisted with data analysis. Yara Hussein (YH) and Wote Amelo Rike (WAR) assisted with cell culture. Idan Rosh (IR) provided cell care. Inna Radzishevsky (IR) provided assistance with the metabolomics experiments. Hanna Mandel offered general experimental support. Julia Ladewig (JL) provided the iPSCs. Tzipora C. Falik Zaccai (TCFZ) assisted during the experimental work. Herman Wolosker (HW) assisted with the design and analysis of the metabolomics data. Shani Stern (SS) (corresponding author) conceptualized and supervised the study, analyzed the data, and drafted the manuscript. All authors reviewed the manuscript.

## Data availability

Correspondence and requests for materials should be addressed to Prof. Shani Stern.

## Materials availability

Correspondence and requests for materials should be addressed to Prof. Shani Stern.

## Code availability

Correspondence and requests for materials should be addressed to Prof. Shani Stern.

